# Running Kinesin-1 shapes the microtubule acetylation gradient

**DOI:** 10.1101/2022.12.01.518806

**Authors:** Mireia Andreu-Carbó, Cornelia Egoldt, Marie-Claire Velluz, Charlotte Aumeier

## Abstract

The properties of single microtubules within the microtubule network can be modulated through posttranslational modifications (PTMs), including acetylation within the lumen of microtubules. To access the lumen, the enzymes could either enter through the microtubule ends or at damage sites along the microtubule shaft. Here we show that the acetylation profile depends on damage sites, which can be caused by the motor protein kinesin-1. Indeed, the entry of the deacetylase HDAC6 into the microtubule lumen depends on kinesin-1-induced damage sites. In contrast, activity of the microtubule acetylase αTAT1 is independent of kinesin-1 and shaft damage. On a cellular level, our results show that microtubule acetylation distributes in an exponential gradient. This gradient results from tight regulation of microtubule (de-)acetylation and scales with the size of the cells. The control of shaft damage represents a novel mechanism to regulate PTM inside the microtubule by giving access to the lumen.

## INTRODUCTION

The molecular motor kinesin-1 transports cargo from the cell center along microtubules towards the cell periphery^1^. To regulate cargo delivery, kinesin-1 activity is tightly regulated. In cells, about 70 % of the kinesin-1 pool is autoinhibited, and kinesin-1 only runs on microtubules when bound to cargo^2,3^. In addition, kinesin-1 runs preferentially on a subset of microtubules within the dense network. This preferential binding can be modulated by Microtubule-Associated Proteins (MAPs), which enhance or inhibit kinesin-1 association to microtubules and impact its running processivity^4–8^.

In addition to MAP-regulated kinesin-1 activity, post-translational modifications (PTMs) of microtubules can increase kinesin-1 binding. In the cellular network, kinesin-1 binds preferentially to a subset of long-lived microtubules which are detyrosinated and/or acetylated^9–14^. Due to its preferential binding, an immotile mutant of kinesin-1 has been proposed as a live-cell marker for acetylated microtubules^15^, although in vitro assays showed that acetylation alone does not increase kinesin-1 binding^16–20^. The strong preferential binding of kinesin-1 to acetylated microtubules in cells is most likely mediated by MAPs. Microtubule acetylation is strongly enriched around the nucleus and decreases towards the cell periphery. Along a single microtubule acetylation is not homogenous; instead, acetylated regions are interspersed with deacetylated regions, resulting in discrete acetylated segments^21–23^.

The α-tubulin acetyltransferase (αTAT1) acetylates microtubules within their lumen on lysine 40 (K40) of α-tubulin^16,24^. In order to acetylate within the lumen, αTAT1 needs to enter the microtubule. The enzyme can enter through the microtubule ends and through damage sites along the microtubule shaft^25–27^. Once inside the microtubule, αTAT1 was suggested to rapidly diffuse and stochastically acetylate tubulin all along the microtubule length^25^. However, a homogenous acetylation along the microtubule is inconsistent with the existence of discrete acetylated segments. The discrete acetylation pattern might result from localized entry of αTAT1 through damage sites and slow diffusion along the microtubule lumen^26,27^. Here we show that the existence of discrete acetylated segments might result from local deacetylation.

The histone deacetylase 6 (HDAC6) removes the acetyl group from α-tubulin^28^. *In vitro* experiments demonstrated that HDAC6 can deacetylate assembled microtubules and tubulin dimers, but deacetylates tubulin dimers 1500-fold more efficiently^28–31^. Microtubules are deacetylated stochastically along their entire length^30,31^. This is consistent with the idea that HDAC6 enters the microtubule lumen in a similar way to αTAT1, although HDAC6 is three-times larger than αTAT1 (140 kDa versus 45 kDa)^28,32^. This difference in the size of their folded domains (Extended Data Fig. 1), could affect their differential ability to enter through damage sites and, once inside, their differential diffusion along the ∼17 nm microtubule lumen.

Damage sites along the microtubule can result from different phenomena: i) imperfect polymerization, ii) occur spontaneously, iii) mechanical forces, or iv) activity of proteins including severing enzymes. Molecular motors do not only use microtubules to transport their cargo, but also generate stress within the microtubule which can damage their underlying microtubule tracks^33–39^. We recently showed that running kinesin-1 enhances tubulin dissociation along the microtubule shaft. This increase in shaft damage depends on the running motion of kinesin-1. Indeed, an immotile kinesin-1 mutant which covers the microtubule did not damage the shaft, but instead reduced shaft damage^37^. Since αTAT1 and HDAC6 entry into the microtubule might depend on damage sites, we investigate here how motor-induced damage sites impact microtubule acetylation.

Here, we report that running kinesin-1 decreases the levels of acetylated microtubules in cells. We show that acetylation levels are decreased around shaft damage sites. In order to manipulate the generation of shaft damage sites, we increased or decreased kinesin-1 activity. Increasing kinesin-1 activity decreases microtubule acetylation, independently of αTAT1 activity. We postulate that, to deacetylate microtubules from within, HDAC6 entry into the lumen depends on damage sites and that kinesin-1 is a potent factor to generate these entry sites.

## RESULTS

### Acetylated microtubule array in cells

The signal of microtubule acetylation was high close to the nucleus and decreased towards the cell periphery (Fig. 1a and see^21–23^). Within the microtubule network of HeLa WT cells, acetylation was discontinuous, and the length of the acetylation segments were on average 2.5 ± 0.71 μm long (Fig. 1b). In the cell center, this results in a dense, but disconnected acetylation array. To quantify the level of acetylation within the microtubule network, we segmented the microtubule network and the acetylation array, and divided the total length of acetylated microtubule segments by the total microtubule length (see Material and Methods). Although the level of acetylated microtubules varied strongly among control cells, on average 36% of the total microtubule network was acetylated in HeLa cells (Fig. 1c). When we measured the fluorescent intensity distribution of acetylation relative to the microtubule network, acetylation exponentially decreased with increasing distance from the cell center. Strikingly, all the different acetylation profiles collapsed onto one master curve when the relative fluorescent intensity and the relative distance are considered (Fig.1d and Extended Data Fig. 2a-c). The characteristic length λ of the exponential fit is 0.3 of the distance from the nucleus to the plasma membrane. This finding implies that acetylation of microtubule scales in cells (Extended Data Fig. 2d).

**Fig. 1.**
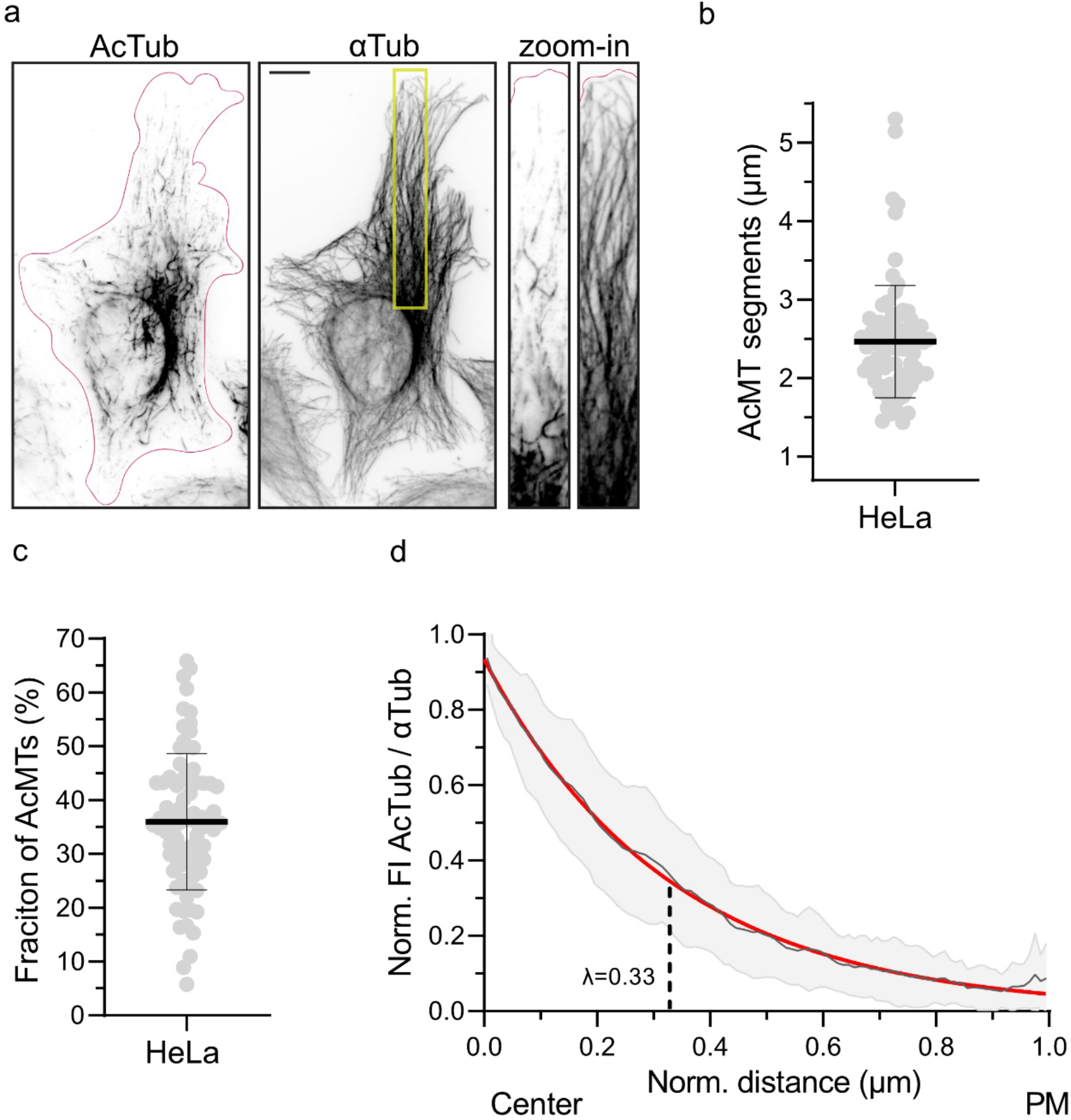
Acetylated microtubule array in HeLa cells. **a**, Representative immunofluorescence images of HeLa cells stained for acetylated tubulin (AcTub) and α-tubulin (αTub). The magenta outline defines the edges of the cell. Scale bar: 10 μm. Right panels show the high magnification of the boxed area for both images. Note the decrease of acetylation levels from the cell center to the cell periphery and the discrete acetylated microtubule segments. **b**, Length of acetylated microtubule segments (AcMT segments) in HeLa cells (n = 80 cells) from 3 independent experiments. Statistics: two tailed t test. Mean with SD. **c**, Fraction of acetylated microtubules (total acetylated microtubule length / total microtubule network length) after skeletonizing the microtubule network in HeLa cells (n = 80 cells) from 3 independent experiments. Statistics: two tailed t test. Mean with SD. **d**, Mean acetylation profile (gray) with exponential fit (red) and the characteristic length λ. Function of the normalized intensity maxima (ActTub/αTub) to the normalized distance from the nucleus (center) to the plasma membrane (PM) in HeLa cells (n= 42 cells) from 3 independent experiments.

### Acetylation is independent from kinesin-1-induced damage sites

In order to study whether αTAT1 entry into the microtubule lumen depends on damage sites, we reduced the number of kinesin-1-induced damage sites by knocking down endogenous kinesin-1 in HeLa cells (see Methods)^36,37^. Western Blot analysis showed that a decrease of kinesin-1 by 85% did not impact the global tubulin acetylation level (Fig. 2a). This is further supported by immunofluorescent analysis, showing a similar level of acetylated microtubules in control and siKinesin-1 transfected cells (Fig. 2b,c). Only the average length of the acetylated segments was slightly longer when kinesin-1 levels were reduced (Fig. 2d). As a reduction of kinesin-1 levels had only a mild effect on microtubule acetylation levels, this suggests that αTAT1 activity is independent of damage sites caused by kinesin-1 activity.

**Fig. 2.**
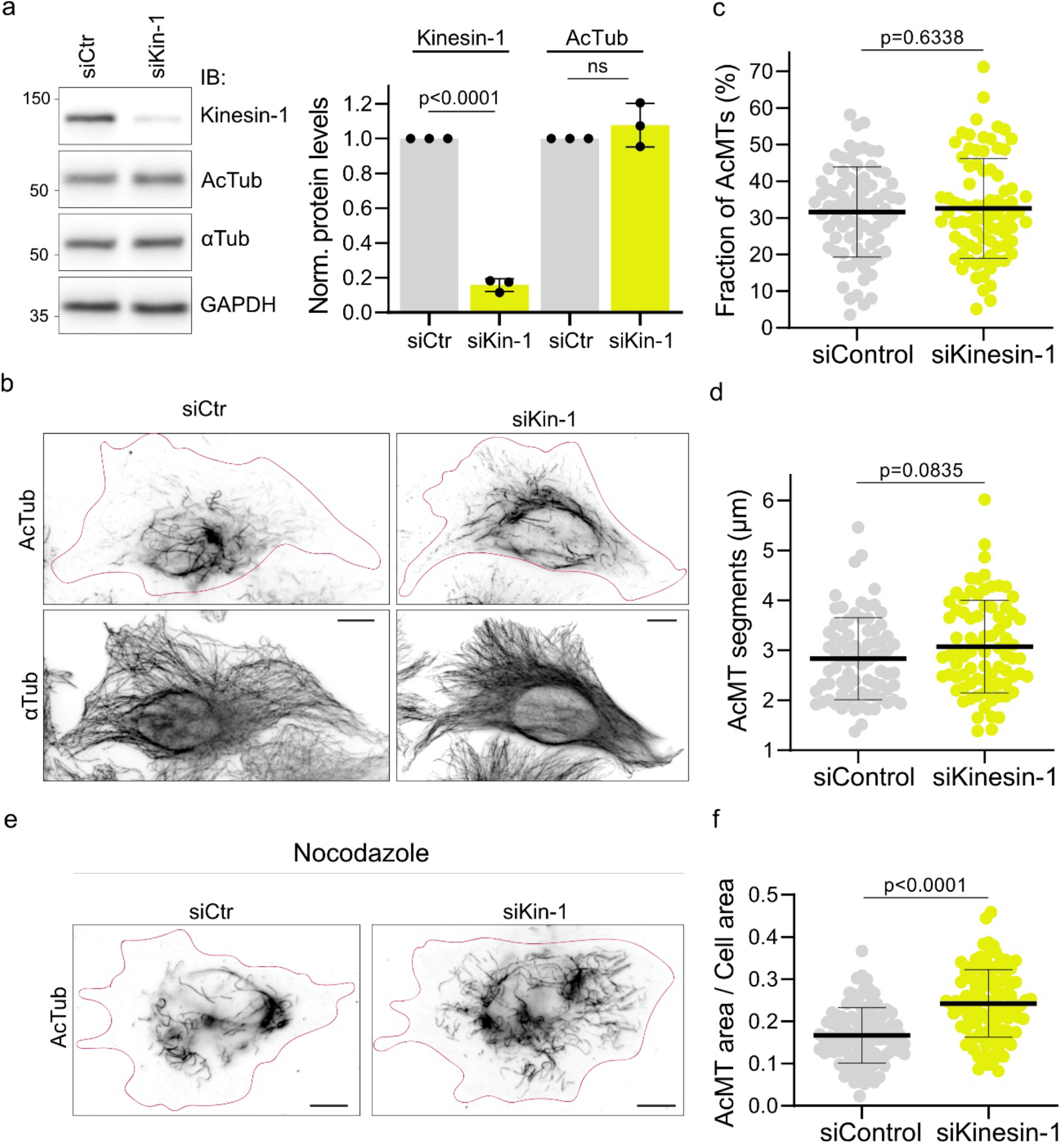
Reduced kinesin-1 activity increases the length of acetylated segments and expands the acetylation array. **a**, Representative western blot analysis with quantification of kinesin-1 and AcTub levels relative to GAPDH in HeLa cells knock-down for kinesin-1 (siKin-1) compared to control cells (siCtr) from 3 independent experiments. Statistics: two tailed t test. Mean with SD. **b**, Representative immunofluorescence images of HeLa siCtr (left) and siKin-1 (right) stained for AcTub and αTub. The magenta outline defines the edges of the cells. Scale bar: 10 μm. **c**, Fraction of acetylated microtubules (total acetylated microtubule length / total microtubule network length) after skeletonizing the network in HeLa cells siKin-1 (n = 101 cells) compared to siCtr cells (n = 100 cells) from 3 independent experiments. Statistics: two tailed t test. Mean with SD. **d**, Length of acetylated microtubule segments in siKin-1 cells (n = 101 cells) compared to siCtr cells (n = 100 cells) from 3 independent experiments. Statistics: two tailed t test. Mean with SD. **e**, Representative immunofluorescence of HeLa kinesin-1 knock-down and control cells treated with 5 μM Nocodazole for 1 h at 37°C before fixation. Cells were stained for AcTub and αTub. The magenta outline defines the edges of the cells. Scale bar: 10 μm. **f**, Area of acetylated microtubule array / total cell area in siKin-1 (n = 112 cells) compared to control cells (n = 101 cells) from 2 independent experiments. Statistics: two tailed t test. Mean with SD.

In cells, a subset of microtubules is long-lived, resistant to microtubule destabilizing drugs (e.g., Nocodazole), and enriched in acetylation^40^. Capitalizing on these properties, we depolymerized dynamic microtubules with nocodazole, to study the impact of reduced kinesin-1 levels on the acetylated microtubule array (Fig. 2e). In control cells, the acetylated array covered 16.7 % of the total area of the cell. Lowering the concentration of endogenous kinesin-1 resulted in a 1.4-fold increase of the acetylated area (Fig. 2f). Consistent with this result, the acetylated array also increased in kinesin-1 knockdown cells in absence of Nocodazole treatment (Extended Data Fig. 3). Therefore, kinesin-1 activity potentially antagonizes microtubule acetylation.

### Running Kinesin-1 reduces the acetylated microtubule array

In different cell lines the endogenous expression levels of kinesin-1 and tubulin acetylation are negatively correlated (Extended Data Fig. 4). To probe to which extend kinesin-1 activity antagonizes microtubule acetylation, we transfected HeLa cells with a construct encoding for a constitutively active, cargo-independent running kinesin-1 (K560-GFP) and immunostained for acetylated tubulin. Expression of running kinesin-1 resulted in a global reduction of acetylation levels (Fig. 3a). In presence of kinesin-1, the acetylation levels decreased up to 2-fold with increasing kinesin-1 concentrations (Fig. 3b). At low K560 expression level, the total acetylation levels were similar to that in control cells (Fig. 3b). However, under these conditions an effect on the spatial organization of the acetylation array was observed. Instead of a dense acetylation cluster at the cell center, short acetylated segments were more dispersed throughout the cell (Fig. 3a,c). Indeed, the average length of acetylated segments was reduced by 20% in cells expressing low levels of kinesin-1 compared to control cells (Fig.3c). With increasing kinesin-1 expression levels, the acetylated segments were even shorter with an average length of 1.2 ± 0.16 μm (compared to 2.5 ± 0.71 μm in control cells) (Fig. 3c). Therefore, running kinesin-1 reduces microtubule acetylation.

**Fig. 3.**
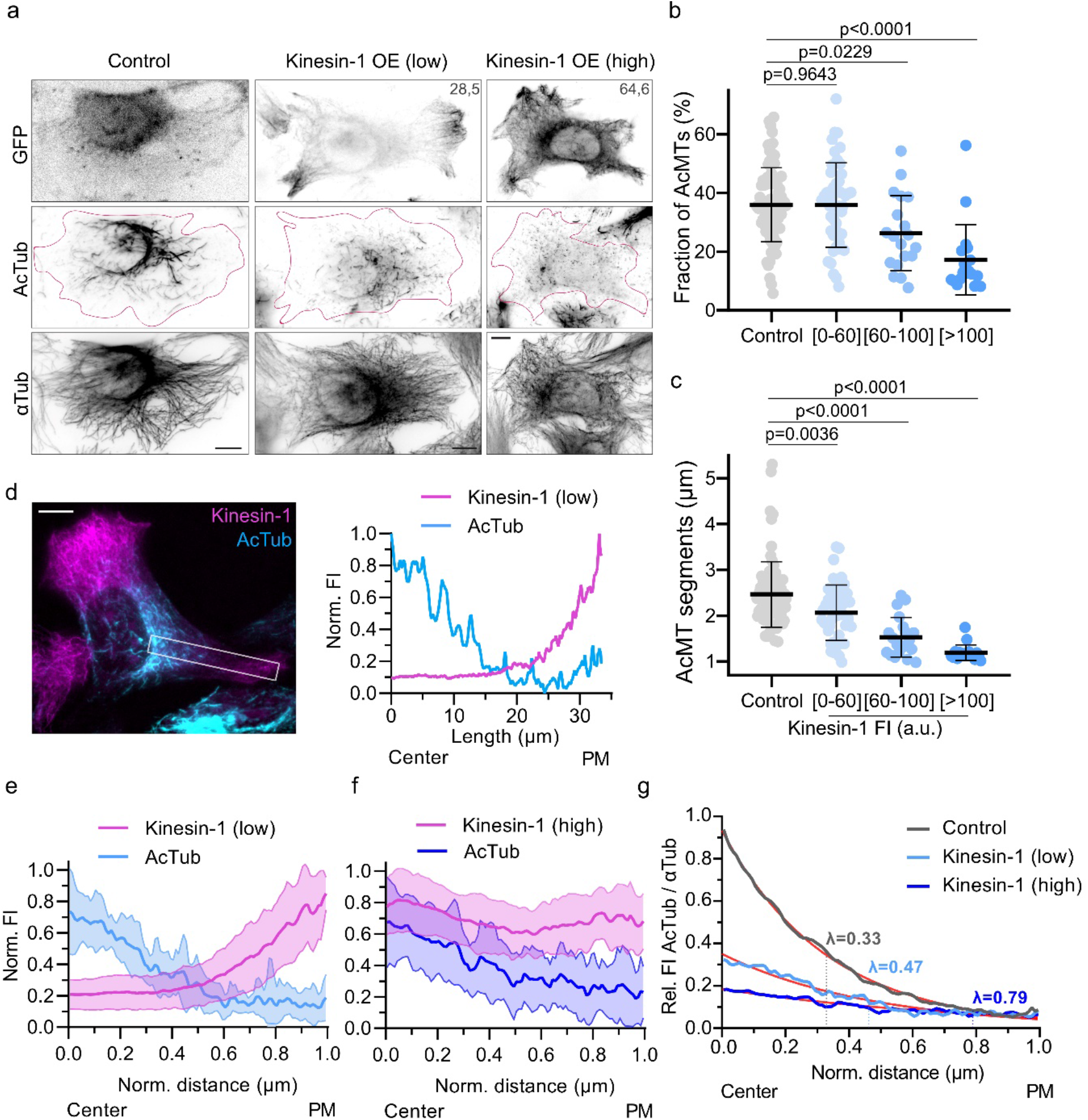
Active kinesin-1 reduces and reorganizes the acetylated microtubule array in cells. **a**, Representative immunofluorescence images of HeLa cells overexpressing pEGFP-N1 (Control) or K560-GFP (kinesin-1 OE) at different expression levels and stained for AcTub and αTub. Numbers in the images correspond to total kinesin-1 FI (a.u.) used for binning in **b** and **c**. The magenta outline defines the edges of the cells. Scale bar: 10 μm. **b** and **c**, Fraction of acetylated microtubules (**b**) and length of acetylated microtubule segments (**c**) in HeLa cells overexpressing pEGFPN-1 (n = 80 cells) and K560-GFP (n= 83 cells) from 3 independent experiments. Statistics: two tailed t test. Mean with SD. FI of kinesin-1-GFP expression was binned in three groups (0-60, 60-100 and above 100 a.u.). **d**, Representative immunofluorescence image of a HeLa WT cell overexpressing K560-GFP and stained for AcTub (left). Normalized FI profile of kinesin-1 and the acetylation distribution along the cell length, in the boxed area of the image (right). Scale bar: 10 μm. **e** and **f**, Mean acetylation profile (blue) and kinesin-1 profile (magenta) with SD. Function of the normalized intensity maxima (ActTub/αTub, or Kininesin-1/aTub) to the normalized distance from the nucleus (center) to the plasma membrane (PM) in HeLa cells from 3 independent experiments. **e**) Low levels of kinesin-1 overexpression (n = 11 cells), **f**) High levels of kinesin-1 overexpression (n = 19 cells). **g**, Relative mean acetylation profiles in presence of different kinesin-1 expression levels with exponential fit (red) and the characteristic length λ. Control HeLa cells from Fig. 1b, low and high kinesin-1 expressing cells from Fig. 3e,f (for details see Methods).

In cells expressing K560, the motor was distributed in a gradient, with increasing levels from the cell center to the cell periphery (Fig. 3d). Kinesin-1 activity induces microtubule deacetylation and therefore forms a counter gradient to microtubule acetylation (decreasing acetylation levels from the cell center to the periphery) (Fig. 3d). With increasing K560 expression levels the shape of the acetylation gradient changed (Fig. 3e,f), the characteristic length of the gradient increased from 0.33 to 0.79 (Fig. 3g). Consistently, with increasing K560 expression levels the amplitude of the acetylation gradient was strongly reduced, up to 5-fold (Fig. 3g). Taken together, running kinesin-1 induces microtubule deacetylation, shortens acetylation segments, and shapes the acetylation gradient.

### Kinesin-1 based deacetylation depends on HDAC6 activity

Microtubules can be deacetylated by HDAC6. To study whether the level of endogenous HDAC6 is a limiting factor for microtubule deacetylation, we overexpressed HDAC6 in HeLa WT cells. Overexpression of HDAC6 did not impact microtubule acetylation (Extended Data Fig. 5). We next studied whether the observed acetylation gradient resulted from HDAC6 activity in the microtubule lumen, by using an HDAC6 specific inhibitor, Tubacin^41^.

Upon inhibition of HDAC6 with Tubacin for 1h, 80% of the microtubule network was acetylated compared to 30% of the network in control conditions (Fig. 4 and see Fig. 1,3 for control cells). This implies that compared to other tubulin deacetylases like Sirt2, HDAC6 is a potent microtubule deacetylase. In the presence of Tubacin, a reduction of the endogenous kinesin-1 expression level had no impact on the acetylation array (Fig. 4a,b). Similarly, overexpression of active kinesin-1 in HDAC6-inhibited cells did not change acetylation levels (Fig. 4c,d). This supports our observation that αTAT1 activity in the microtubule lumen does not require the presence of damage sites generated by running kinesin-1 (Fig. 2). This implies that i) αTAT1 activity is uncoupled from kinesin-1 activity, ii) Sirt2, another tubulin deacetylase^42^, plays a minor role in microtubule deacetylation, iii) kinesin-1 effects on deacetylation are mediated by HDAC6, and iv) damage sites generated by the running of kinesin-1 are entry points that give HDAC6 access to the lumen.

**Fig. 4.**
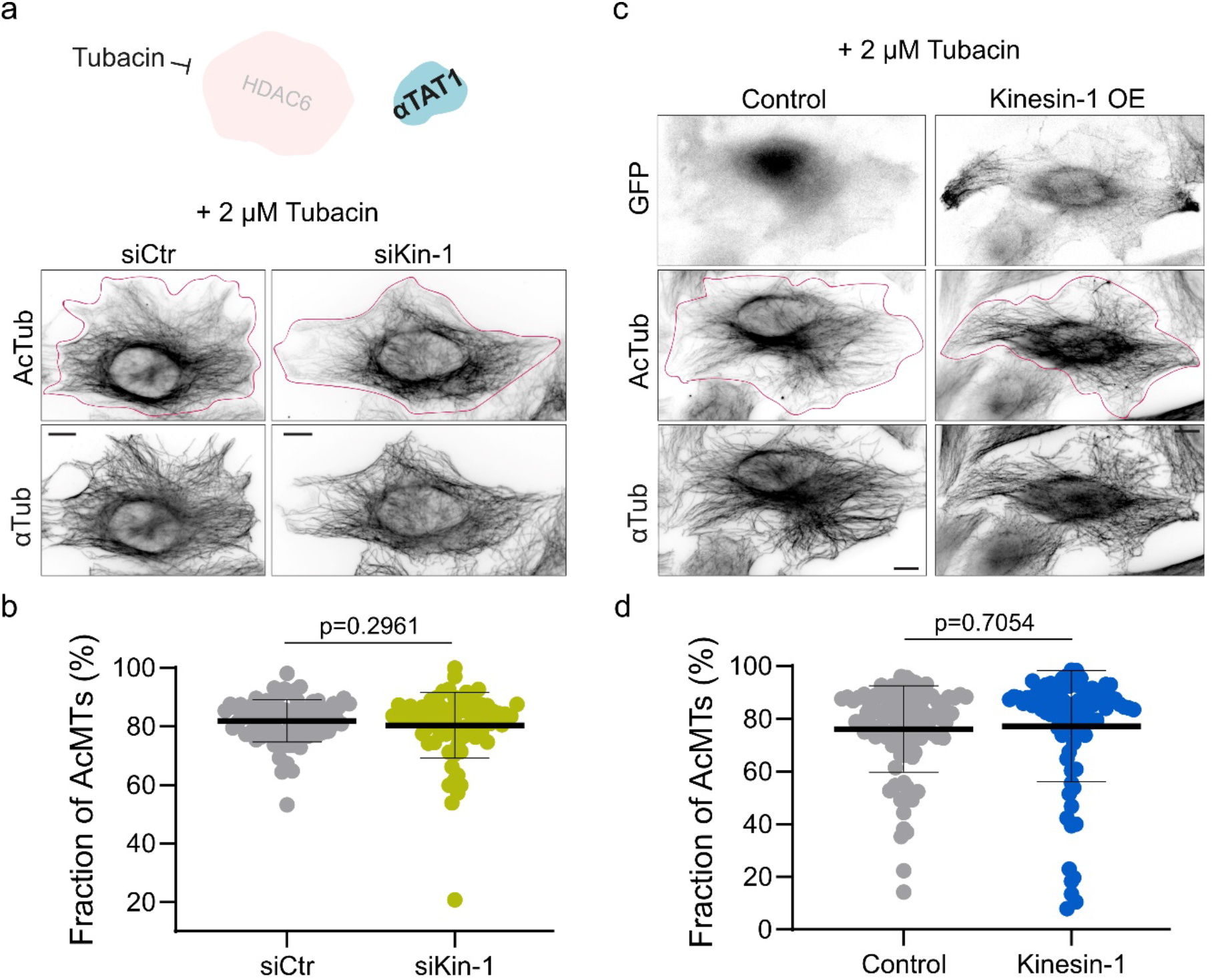
Kinesin-1 based microtubule deacetylation depends on HDAC6, while αTAT1 efficiency is independent of kinesin-1-induced damages. **a**, Representative immunofluorescence of HeLa kinesin-1 knock-down (siKin-1) and control cells (siCtr), treated with 2 μM Tubacin for 1 h at 37°C before fixation. Cells were stained for AcTub and αTub. The magenta outline defines the edges of the cells. Scale bar: 10 μm. **b**, Fraction of acetylated microtubules in HeLa cells siKinesin-1 (n = 75 cells) compared to siControl cells (n = 80 cells) from 3 independent experiments. Statistics: two tailed t test. Mean with SD. **c**, Representative immunofluorescence images of HeLa cells overexpressing pEGFP-N1 (Control) or K560-GFP (Kinesin-1 OE) treated and stained as in (a). Scale bar: 10 μm. **d**, Fraction of acetylated microtubules in HeLa cells overexpressing pEGFPN-1 (n = 89 cells) and K560-GFP (n = 89 cells) from 3 independent experiments. Statistics: two tailed t test. Mean with SD.

### Only running kinesin-1 reduces microtubule acetylation

Running molecular motors can damage the shaft, while immotile kinesin-1 cover microtubules and hinder instead the generation of damage sites^35,37^. We studied whether microtubule deacetylation is mediated by damage sites generated by the running motion of kinesin-1 on microtubules. To do so, we compared microtubule acetylation in HeLa cells expressing a running kinesin-1 versus an immobile kinesin-1 (K560-rigor-GFP)^43,44^. Unlike running kinesin-1, we observed a strong correlation between microtubules which were densely covered by the rigor kinesin-1 and microtubules which were highly acetylated (Fig. 5a). Overexpression of the rigor mutant increased the acetylation levels within the microtubule network by 2.2-fold, rather than reducing them (Fig. 5b). Therefore, kinesin-1 binding to microtubules itself does not explain the kinesin-1-dependent microtubule deacetylation, but like in the case of the generation of the damage sites deacetylation requires the running motor. Because both, the generation of damage sites and the effects on deacetylation, require kinesin motility, this implies that deacetylation itself requires the presence of damage sites.

**Fig. 5.**
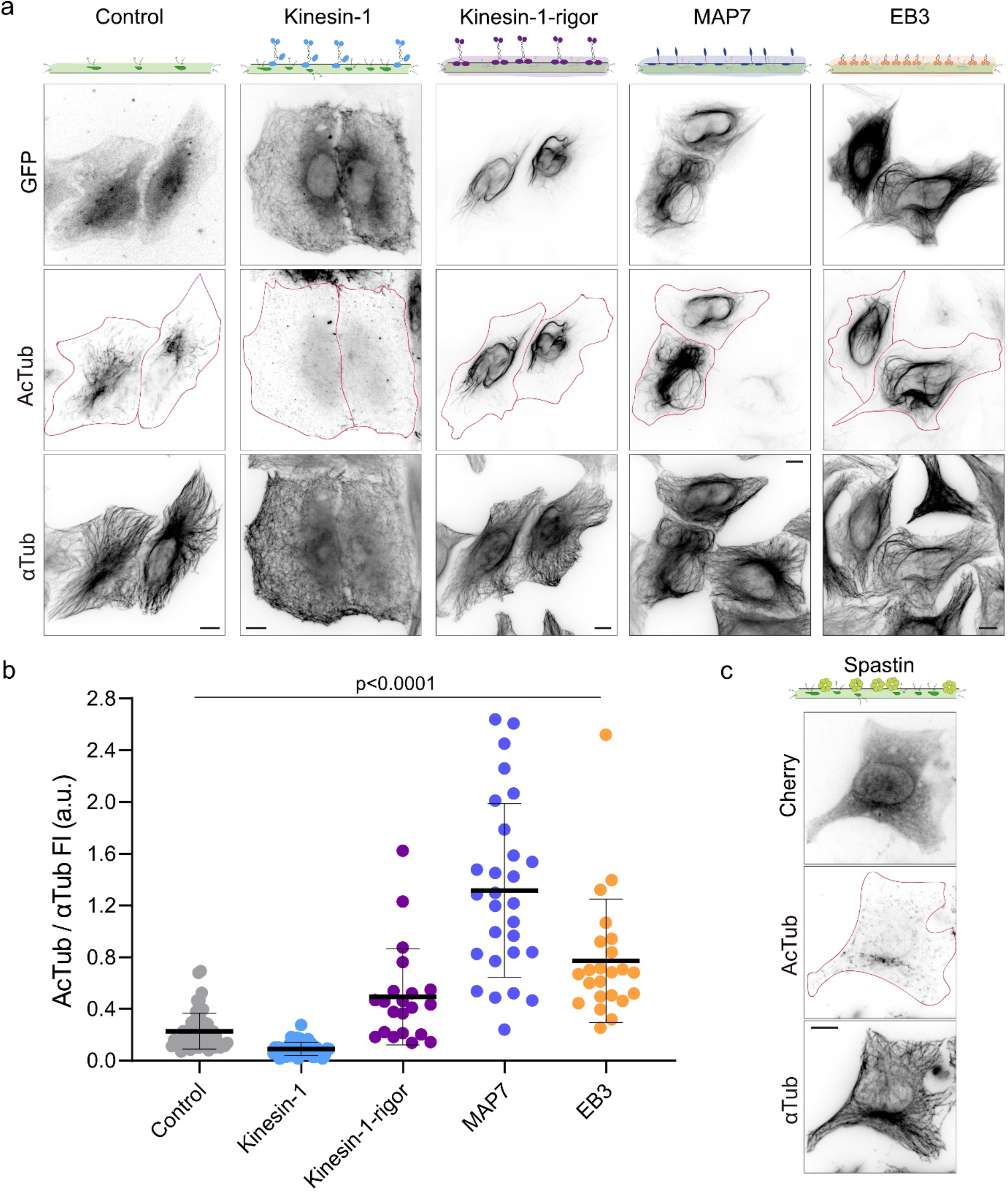
Kinesin-independent shaft damage affects microtubule deacetylation. **a**, Representative immunofluorescence images of HeLa cells overexpressing pEGFP-N1 (Control), K560-GFP (kinesin-1), K560-rigor-GFP (kinesin-1-rigor), MAP7-GFP (MAP7) or EB3-GFP (EB3) and stained for AcTub and αTub. The magenta outline defines the edges of the overexpressing cells. **b**, FI of the AcTub relative to the total FI of αTub per cell in each condition in **a**. Note that values above 1 are due to differences in the acetylation and tubulin antibodies and are not informative for the fraction of AcTub (for details see Methods). Statistics: two tailed t test. Mean with SD from 2-3 independent experiments. **c**, Representative immunofluorescence images of HeLa cells overexpressing mCherry-Spastin (Spastin) and stained for AcTub and αTub. The magenta outline defines the edges of the cell. Scale bar: 10 μm.

### MAPs covering the microtubule shaft increase microtubule acetylation

We next asked whether increasing the acetylation level is specific for the kinesin-1 rigor mutant, or a general property of MAPs covering the microtubule shaft. To address this, we expressed in HeLa cells the microtubule-associated protein 7 (MAP7) and the end-binding protein 3 (EB3). When overexpressed in cells, both MAPs decorated the microtubule shaft (Fig. 5a)^6,45^. Remarkably, overexpression of MAP7 and EB3 caused hyperacetylation of the microtubule network compared to non-transfected neighboring cells (Fig. 5a, magenta outlines). Of all three proteins, MAP7 showed the highest level of microtubule acetylation, with a 6-fold increase compared to control cells (Fig. 5b). Taken together, these results uncover a scenario where proteins covering the microtubule shaft increase acetylation possibly by hindering the formation of damage sites. In this context, the effect of kinesin-1 on deacetylation differs from other MAP proteins, because the running of the motor itself causes the generation of damage sites.

### Shaft damage independent of kinesin deacetylates microtubules

Shaft damage by itself, beyond damage generated by running kinesin-1 in particular, could be a general mechanism to deacetylate microtubules. To address this, we damaged microtubules using severing enzymes^*46*^ instead of kinesin-1. We transfected HeLa cells with mCherry-Spastin and immunostained for acetylated tubulin. As previously reported, high levels of Spastin overexpression severed the microtubule network into fragments (Extended Data Fig. 6). At low levels of Spastin expression, cells had an intact microtubule network, but the shaft might be damaged by incomplete severing activity (Fig. 5c). These cells hardly showed any acetylated microtubules, consistent with the idea that low levels of Spastin boost the generation of damage sites ^*46*^, which could give access to HDAC6 and thereby boost deacetylation. Consistently, the remnants of acetylation segments are reduced to short foci (Fig. 5c). Therefore, the control of shaft damage sites beyond those generated by kinesin-1 is a general mechanism by which microtubules could be deacetylated.

### Microtubules are deacetylated around damage sites

To study whether microtubule deacetylation occurs at damage sites, we analyzed the local acetylation levels around the damage sites by using a damage/repair site-specific antibody^47,48^ and co-stained for acetylation. In WT cells the levels of damage/repair sites increased from the cell center to the periphery (Fig. 6a). Consistent with our observation that K560 induces microtubule deacetylation close to the cell center, in presence of K560 the distribution of damage/repair sites increased at the cell center, leading to a more homogenous distribution of damage/repair sites throughout the cell (Fig. 6c,d).

**Fig. 6.**
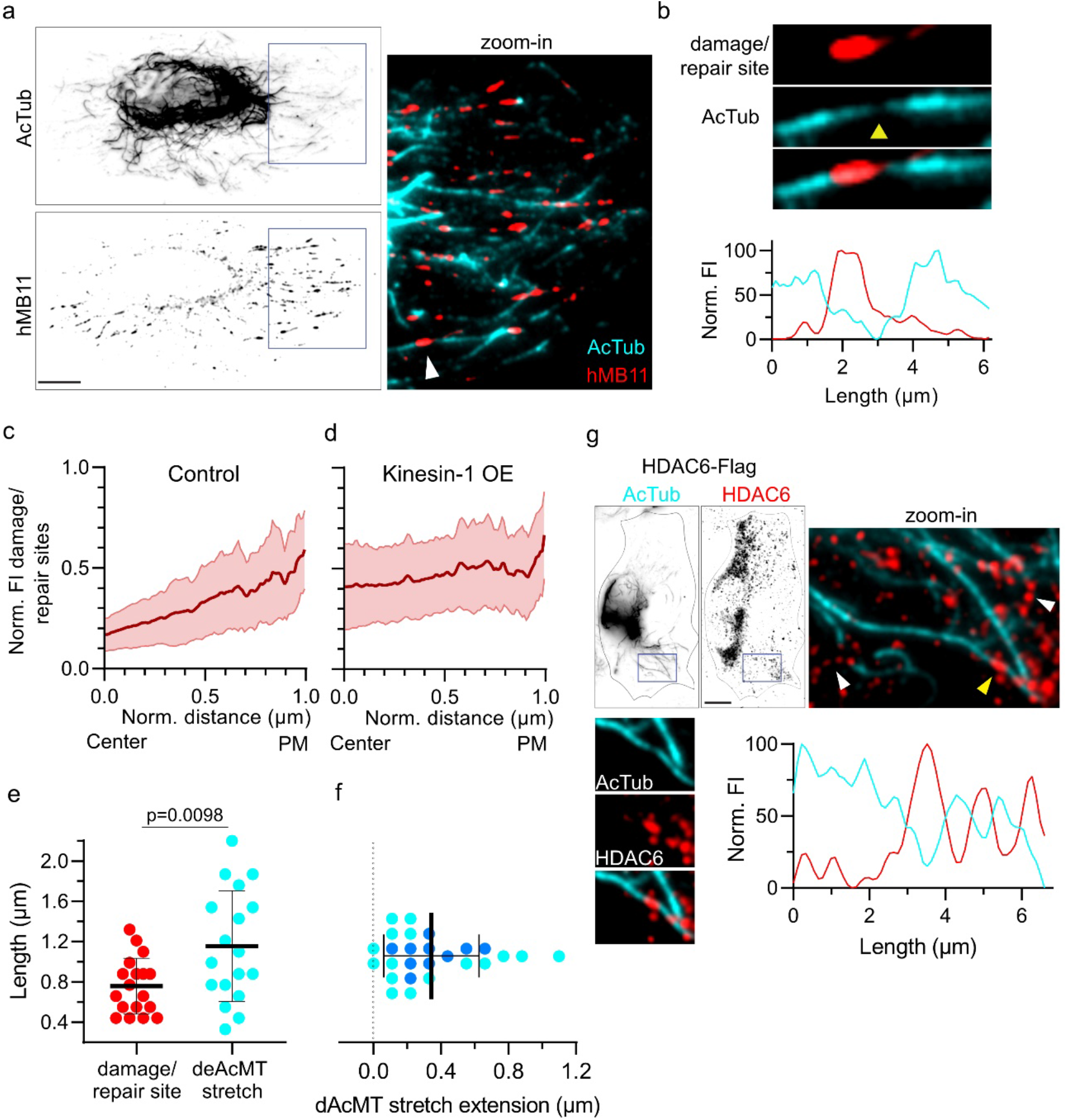
Microtubules are deacetylated at damage/repair sites. **a**, Representative immunofluorescence image of a HeLa WT cell stained for damage/repair sites (hMB11, also labeling the growing microtubule tip) and AcTub. Scale bar: 10 μm. The zoom-in shows the merge of the two images, white arrowhead points to an acetylated microtubule with a damage/repair site. **b**, Representative acetylated microtubule with a damage/repair site. Yellow arrowhead indicates deacetylation stretch; corresponding line scan below. **c** and **d**, Spatial distribution profile of the hMB11 FI relative to the FI of αTub, normalized intensity maxima, in cells overexpressing pEGFPN-1 (Control, n = 45, **c**) and kinesin-1 overexpression (kinesin-1 OE, n = 40, **d**) from 3 different experiments. Red lines: Mean with SD. **e**, Length of damage/repair sites (red) and deacetylated stretches (cyan). Mean with SD of a total of n = 18 sites from 1 experiment. Statistics: two tailed t test. **f**, Extension of deacetylate stretch beyond repair/damage sites. Light blue dots correspond to a microtubule plus-end directed extension and dark blue dots correspond to a minus-end directed extension with respect to the damage/repair site (0.0 = border of damage/repair site). **g**, Representative immunofluorescence image of a HeLa WT cell overexpressing HDAC6-Flag and stained for AcTub and Flag. Scale bar: 10 μm. Zoom-in of the merge of the two images in **a** (right). White arrowheads point to regions where HDAC6 colocalizes with deacetylated microtubules. Yellow arrowhead points to an acetylated microtubule showing HDAC6 colocalization; below corresponding zoom-in with split channels (right) and the line scan (below).

Figure 6a,b shows that microtubule damage sites are embedded within stretches of deacetylated microtubules. The deacetylated stretches were 1.5-fold longer than the damage sites (Fig. 6e) and did extend 0.34 μm beyond the damage sites (Fig. 6f). At the border of the deacetylation stretch the acetylation level increased gradually. Therefore, microtubule deacetylation cannot only result from the repair of the damage sites with non-acetylated tubulin, but implies the entry of HDAC6 at damage sites and deacetylation activity around damage sites. Indeed, HDAC6 can be seen as associated to these stretches of deacetylated microtubules (Fig. 6g). Taken together, the association of non-acetylated tubulin around damage sites, the gradual profile of deacetylation and the spatial association of HDAC6 to these phenomena, uncover a scenario in which the damage sites are entry points for HDAC6 into the microtubule lumen. Once inside, this deacetylase can undergo limited diffusion, probably due to its large size, generating local islands of deacetylation.

## DISCUSSION

Microtubules are acetylated within their lumen. To modify the microtubule inside, the enzyme responsible for tubulin acetylation, αTAT1, needs access to the lumen. We show that running kinesin-1 does not affect microtubule acetylation, but causes deacetylation mediated by HDAC6. Considering our previous results showing that active, running kinesin-1 by itself induces shaft damage^37^, we propose a model for the control of the acetylation state of microtubules in which i) the smaller αTAT1 acetylation enzyme can rapidly diffuse throughout the lumen after entry from the microtubule tips^25^, ii) while the deacetylation enzyme HDAC6, which is threefold larger than αTAT1 and diffuses poorly, critically depends on damage sites along the shaft to access the microtubule lumen, and iii) once in the lumen HADC6 deacetylates stretches around the damage sites. Poor HDAC6 diffusion within microtubules is consistent with the fact that its deacetylation activity in intact microtubules is much lower than its activity on non-polymerized tubulin dimers or dimers polymerized in open sheets^31^.

Shaft damage can regulate microtubule length and lifetime^36,37,46,49^. In addition, our data show that damage sites can also regulate microtubule posttranslational modifications (Fig. 6c-g). Therefore, a control handle on the generation of damage sites represents a regulatory mechanism for microtubule network organization and heterogeneity. The effect of the severing protein Spastin, as well as of the MAPs EB3 and MAP7 on the generation of damage sites and the acetylation state shows that not only motors, but a palette of different microtubule binding factors can contribute to the regulation of microtubule shaft dynamics.

Although many studies address microtubule acetylation, it was still unknown what establishes the acetylation pattern in cells. We show that microtubule acetylation is distributed in a gradient. A cellular gradient opens the possibility of positional information. It will be of future interest to study which proteins can “read” the acetylation gradient. Furthermore, what scales the acetylation gradient? Surprisingly, the running activity of kinesin-1 is one parameter that can shape the acetylation gradient, by damaging the microtubule shaft. Generation of damage sites by different proteins might be a general mechanism to scale the gradient, which implies a tight regulation of shaft integrity. However, further factors might be involved in establishing the acetylation gradient, including different effective acetylation and deacetylation rates, or defined spatial lumen entry for αTAT1 and HDAC6.

## ACKNOWLEDGEMENTS

We thank M. Gonzalez-Gaitan and R. Wimbish for careful reading of our manuscript.

## CONTRIBUTIONS

MAC and CA conceptualized the study. MAC and CA designed the experiments. MAC performed and analyzed all the experiments with the help of CE. MCV cloned all the plasmids used in this study.MAC and CA wrote the manuscript.

## METHODS

### Cell culture, transfection and siRNA knockdown

HeLa (ATTC-CCL-2™) and U-2 OS (a gift from Laurent Blanchoin) cells were cultured in high glucose Dulbecco’s Modified Eagle’s Medium (DMEM, ThermoFisher, 61965026), hTETR-RPE1 (ATCC-CRL-4000) cells were cultured in high glucose Dulbecco’s Modified Eagle’s Medium F12 (DMEM, ThermoFisher, 113057), and supplemented with 10 % Fetal Bovine Serum (FBS, ThermoFisher, 10270106) and 1 % penicillin-streptomycin (Gibco, 15140122). Ptk2 (ATCC-CCL-56) cell line was grown in Minimum Essential Medium Eagle (MEM, Sigma, M0643) supplemented with 10 % Fetal Bovine Serum, 1 % penicillin-streptomycin and 1 % L-Glutamine (Gibco, 25030024). All cell lines were kept at 37°C with 5 % CO_2_ and monthly checked for mycoplasma contamination.

For transient protein overexpression, HeLa cells were grown until 60-70 % confluency in 6-well plates (on 12 mm coverslips [3 to 4 per well] for immunofluorescence experiments). HeLa cells were transfected with jetOPTIMUS transfection reagent (Polyplus) using 1 μg of plasmid in 6-well plates according to the manufacturer’s instructions. 4 h post-transfection, the transfection media was replaced with fresh media. Cells were analyzed 18-24 h after transfection. The constructs of human constitutively active form of kinesin-1 (K560-GFP) and the ATPase rigor mutant of K560 (K560-rigor-GFP) were transfected as previously described^37^. The microtubule-binding domain of enconsin (18-283 amino acids) (addgene #26741), referred in the text as MAP7, was subcloned into pEGFP-C1 using EcoRI/AgeI restriction enzymes. The mCherry-EB3-7 vector was obtained from Addgene (addgene #55037) and the mCherry was removed and replaced by a GFP sequence using AgeI/BsrG1 restriction enzymes. The plasmid encoding Drosophila melanogaster full-length Spastin was a kindly gift from Dr. Antonina Roll-Mecak and cloned into pmCherry-C1 vector (mCherry-Spastin) using EcoRI/KpnI restriction enzymes. In control experiments, the empty vectors pEGFP-N1 or pmCherry-C1 were transfected.

For generating kinesin-1 knockdown cells, HeLa cells were transfected with a combination of four siRNA duplexes against kinesin-1 (Kif5B) as previously described^37^.

### Drug treatment

To depolymerize the dynamic microtubule network, HeLa cells were treated with 5 μM Nocodazole (Sigma, M1404) diluted in culture media for 1 h at 37°C prior to fixation. To hyperacetylate the microtubule network, we used the specific HDAC6 inhibitor Tubacin^41^. HeLa cells were treated with 2 μM Tubacin (Sigma, SML0065) diluted in culture media for 1 h at 37°C before fixation. In control experiments, DMSO was added at the same volume as the drugs.

### Immunofluorescence

18-24 h after transfection, HeLa cells were fixed with 100 % cold methanol for 4 min at −20°C. Cells were then permeabilized and blocked with the blocking buffer [0.1% Triton X-100 and 2% bovine serum albumin in phosphate buffered saline (PBS)] for 30 min and subsequently incubated with the primary antibodies rabbit anti-αTubulin (Abcam, ab18251,1:1000 dilution), mouse anti-acetylated Tubulin (Sigma, T74451, 1:1000 dilution) or rabbit anti-Flag (Sigma, F7425, 1:500 dilution) in blocking buffer in an humidified chamber at room temperature for 1 h. Cells were washed in PBS three times for 10 min at room temperature, then subsequently incubated in secondary antibodies (Invitrogen, species-specific IgG conjugated to Alexa-647, 561, or 488 fluorophores, 1:1000 dilution in blocking solution) for 1 more h in an humidified chamber at room temperature. Finally, cells were washed three more times with PBS, and coverslips were mounted onto glass microscopy slides (Glass technology) using ProLong™ Diamont Antifade Mountant.

To visualize shaft damages, we used the damage/repair specific antibody hMB11 as previously described^37^ before methanol fixation and the immunofluorescence procedure explained above.

Immunofluorescence images were acquired using an Axio Observer Inverted TIRF microscope (Zeiss, 3i) and a Prime BSI (Photometrics) using a 100X objective (Zeiss, Plan-Apochromat 100X/1.46 oil DIC (UV) VIS-IR). SlideBook 6 X64 software (version 6.0.17) was used for image acquisition.

### SDS-PAGE and Western blot

Cells were lysed [50 mM Tris-HCl (pH 7.5), 150 mM NaCl, 1 % Triton X-100, 0.5 % SDS, and a protease inhibitor tablet (Roche)] for immunoblotting. Proteins were separated by 8% acrylamide SDS-PAGE gels and then transferred to a nitrocellulose membrane with an iBLOT 2 Gel Transfer Device (ThermoFisher Scientific, IB21001). Nitrocellulose membranes were blocked for 1 h with the blocking buffer containing 5 % dried milk in TBS-Tween 1 % and incubated over-night with primary antibodies: anti-αTubulin (Sigma, T6074, 1:1000 dilution), anti-acetylated Tubulin (Sigma, T74451, 1:1000 dilution) anti-UKHC (Santa Cruz Biotechnology, SC-133184, 1:1000 dilution) and anti-GAPDH (Millipore, MAB374, 1:1000 dilution) in blocking buffer. The next day, membranes were washed three times with TBS-Tween 1 % and incubated for 1 h at room temperature with a secondary antibody anti-Mouse conjugated to horseradish peroxidase (GE Healthcare, NA9310, 1:5000 dilution in blocking buffer), washed three times with TBS-Tween 1 % solution and revealed with an ECL Western blotting detection kit (Advansta) and with Fusion Solo Vilber Lourmat camera (Witec ag).

### Image and statistical analysis

Immunofluorescence images were analyzed using ImageJ. For data analysis we subtracted the background individually for each channels.

To measure the <u>level of acetylated microtubules in cells</u>, individual segments for the acetylated tubulin and α-tubulin channel were detected using CurveTrace plugin v.0.3.5 plugin for ImageJ (https://github.com/ekatrukha/CurveTrace). In a next step, we binarized images of the detected segments for both the acetylated array and microtubule network. The region of interest (ROI) defining single cells was manually drawn using the GFP channel. The total length of all acetylated segments and the total microtubule network length per cell were measured from the binarized images. To quantify the <u>fraction of acetylated microtubules</u>, the total length of acetylated microtubules was divided by the total length of the microtubule network per cell. The <u>length of the acetylated segments</u> represents the average lengths of all acetylated segments in one cell. In addition, in cells overexpressing K560-GFP or pEGFP-N1 the fluorescence intensity of the GFP signal per cell was measured. To measure the <u>area of acetylated microtubules after Nocodazole treatment</u>, we made an acetylation mask based on the fluorescent using the default thresholding process in Image J. The fluorescence intensity of the cytoplasmic tubulin was used to define the area of the cell. Finally, the area of the acetylated array mask was measured and divided by the total area of the cell. <u>Differences in the levels of acetylation</u> in cells overexpressing different GFP-constructs (Fig. 5 and Extended Data Fig. 5,6) were determined by measuring the fluorescence intensity of the acetylated tubulin antibody relative to the fluorescence intensity of the microtubule network with an α-tubulin antibody. Since the antibodies have different sensitivities, values above 1 do not indicate that the acetylated array spanned over the microtubule network. Instead, the acetylated tubulin antibody exhibits higher fluorescent intensity values than the α-tubulin antibody.

For measurements of the <u>distribution of the acetylated microtubule</u>, a straight line of 45-pixel width was manually drawn from the perinuclear region (with the maximum fluorescent intensity) following the microtubule network to the cell periphery. The intensity profiles for each channel were measured. Additionally, the average background fluorescent intensity close to microtubules was measured for each individual channel and subtracted from the measured intensity values. For the acetylation profile, the intensity values for acetylated tubulin were divided by the signal intensities of the α-tubulin line scan. These relative values (Y-axes) and the length of each line scan (X-axes) were divided by the maximum value, in order to normalize each data set. To compare different data sets the X-axes values were binned into 0.005 steps. Resulting acetylation intensity profiles were averaged and fitted to an exponential decay function by setting the minimum Y-axis value to 0 (plateau) to determine the characteristic length λ (nonlinear regression, one phase decay in GraphPad Prism v.9; constraint: set plateau to 0) (Fig. 1b). Kinesin-1 gradients were obtained as described for the acetylation profile here above, but without a plateau constraint. Kinesin-1 overexpression was considered low or high when fluorescence intensity values were lower or higher than 60 a.u., respectively (Fig. 3e,f). In order to compare changes in acetylation decay length upon kinesin-1 overexpression, the amplitude of the control curve was set to 1. The average of the exponential decay curves for low and high kinesin-1 overexpression were plotted as relative of the control, whereas the amplitude (Y_0_) for each condition was determined by the ratio of the fluorescence intensity of acetylated tubulin over α-tubulin at the nucleus (Fig. 3g).

To quantify the <u>length of damage/repair sited and the deacetylated microtubule stretches (dAcMT stretches)</u>, areas with damage repair sites surrounded by acetylation segments were analyzed using the segmented-line tool in ImageJ to obtain the fluorescence intensity profile. Line scans for both channels were normalized and the intensity values above or below 50 a.u. were considered as damage/repair sites or dAcMT stretches, respectively (see Fig. 6b for an example). From those intensity values above or below 50 a.u., we determined the length of damage/repair sites or dAcMT stretches, respectively. For quantifying the damage/repair sites distribution for kinesin-1 overexpression and control cells, line scans were measured and analyzed as described above.

Statistical analyses were performed by two-tailed unpaired Student’s *t* test using GraphPad Prism software v.9. P values less than 0.05 were considered statistically significant.

## Extended data

**Extended Data Figure 1.**
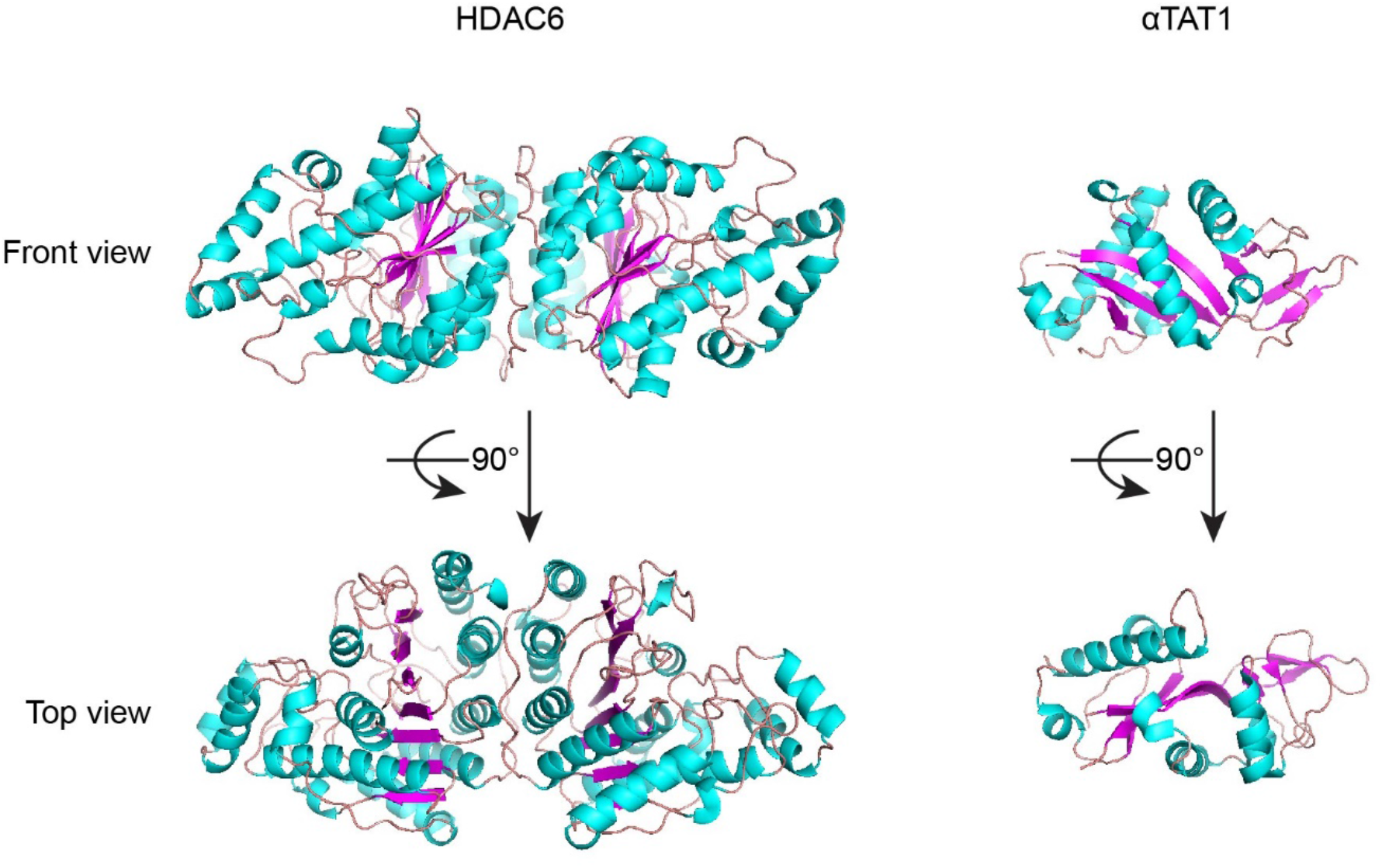
Representation of the structured domains of HDAC6 and αTAT1. Front and top view of the AlphaFold prediction of the structured HDAC6 catalytic domain (AF-Q9UBN7) (left). Front and top view of the αTAT1 structured region (PDB, 4GS4) (right). α-helices are colored in blue and β-strands in pink. Note that both proteins have predicted unstructured domains which are not represented here. Structures were adapted using Pymol.

**Extended Data Figure 2.**
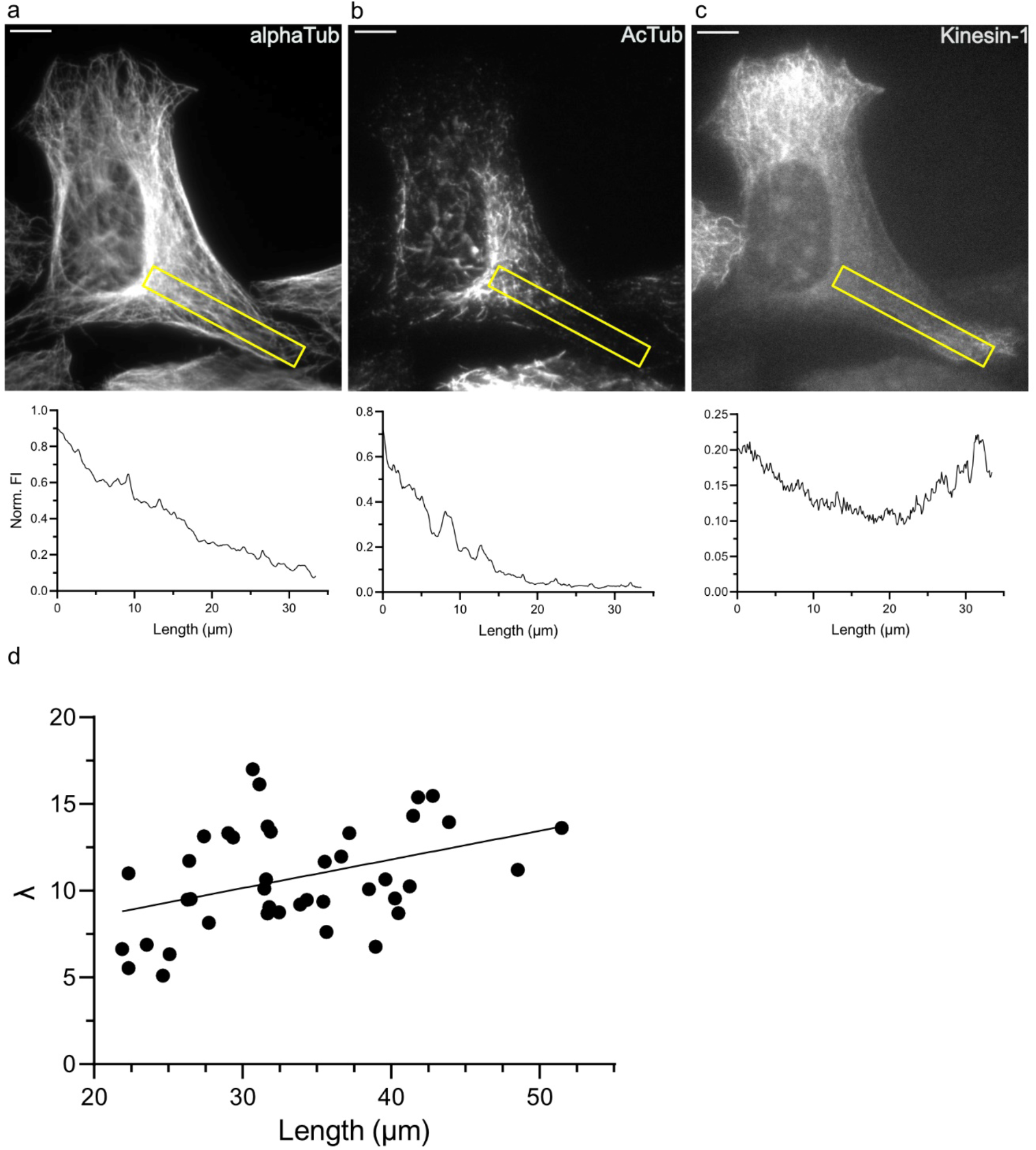
Gradient of acetylated microtubule array in HeLa cells. **a, b** and **c**, Representative cell with selected 45-pixel line for analyses of the fluorescence intensity profile of the microtubule network with corresponding intensity profile normalized to the maximum intensity value (a), acetylated tubulin array (b) and kinesin-1 overexpression (c). Scale bar: 10 μm. **d**, Correlation of the characteristic length λ and the cell length for Control HeLa cells from Fig. 1b. Data were fitted using a simple linear regression (R^2^ = 0.16) using GraphPad Prism software v.9.

**Extended Data Figure 3.**
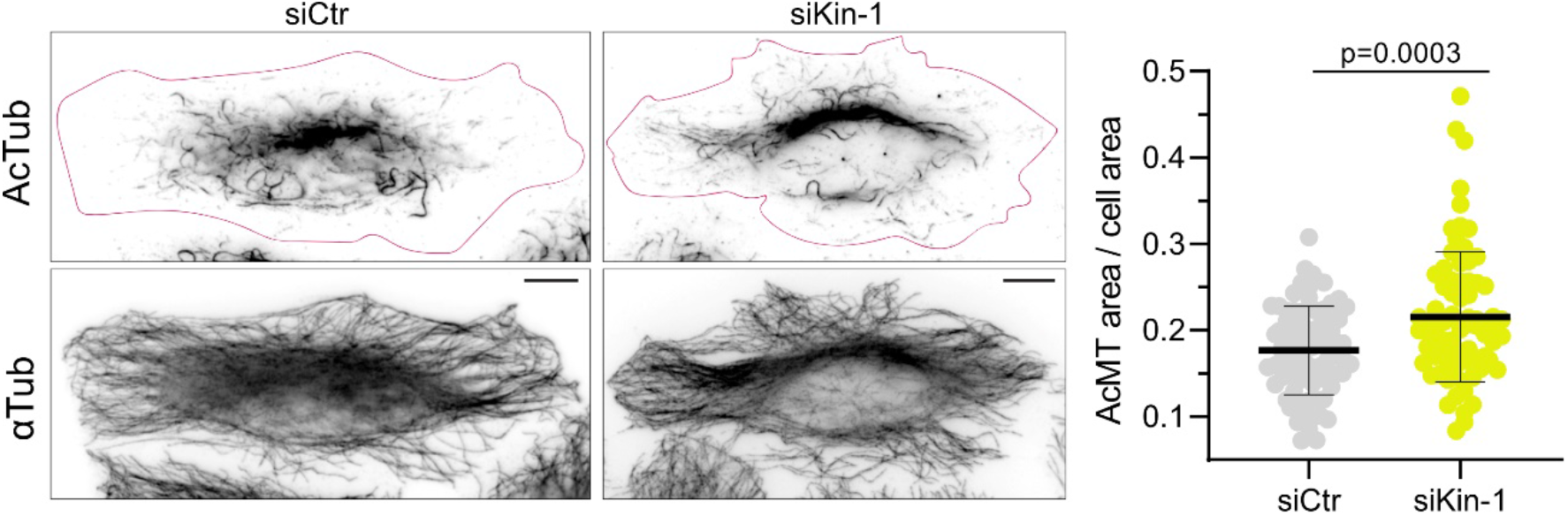
Kinesin-1 knock-down cells display a sparser microtubule acetylated array. Representative immunofluorescence of HeLa kinesin-1 knock-down and control cells and the quantification of the area of acetylated microtubule array / total cell area in siKin-1 (n = 76 cells) compared to control cells (n = 76 cells) from 3 independent experiments. Statistics: two tailed t test. Mean with SD. Cells were stained for AcTub and α-tubulin. The magenta outline defines the edges of the cells. Scale bar: 10 μm.

**Extended Data Figure 4.**
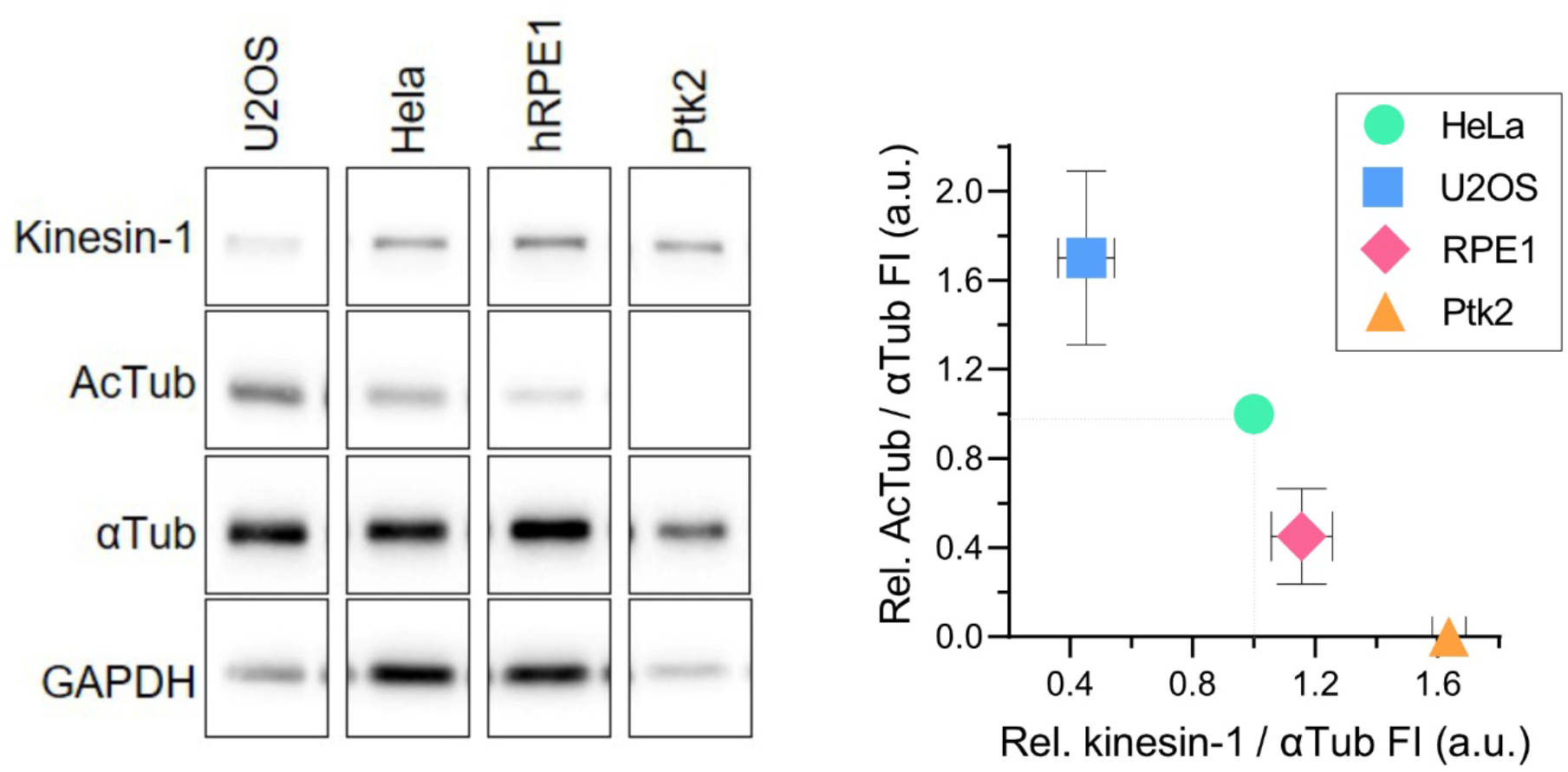
Kinesin-1 and acetylated tubulin levels in different cell lines. Representative western blot (left) with quantification of the levels of kinesin-1 and AcTub relative to αTub (right) in different cell lines from 2 to 4 independent experiments. Relative kinesin-1 and AcTub levels were normalized to the Hela cells values. Mean with SD.

**Extended Data Figure 5.**
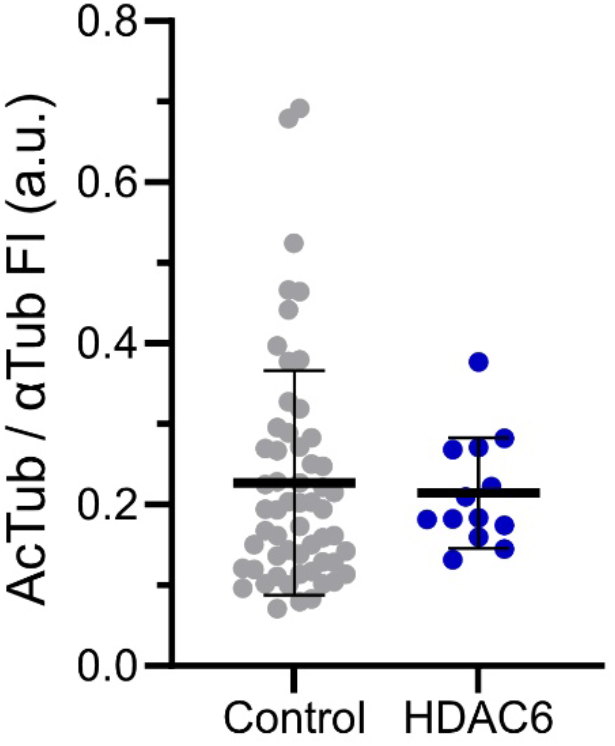
HDAC6 overexpression does not reduce the acetylated microtubule array. Quantification of the AcTub FI relative to the FI of the microtubule network (αTub) in HeLa WT cells overexpressing HDAC6 (n = 13 cells) compared to control cells (n = 30 cells) from 2 independent experiments. Statistics: two tailed t test. Mean with SD.

**Extended Data Figure 6.**
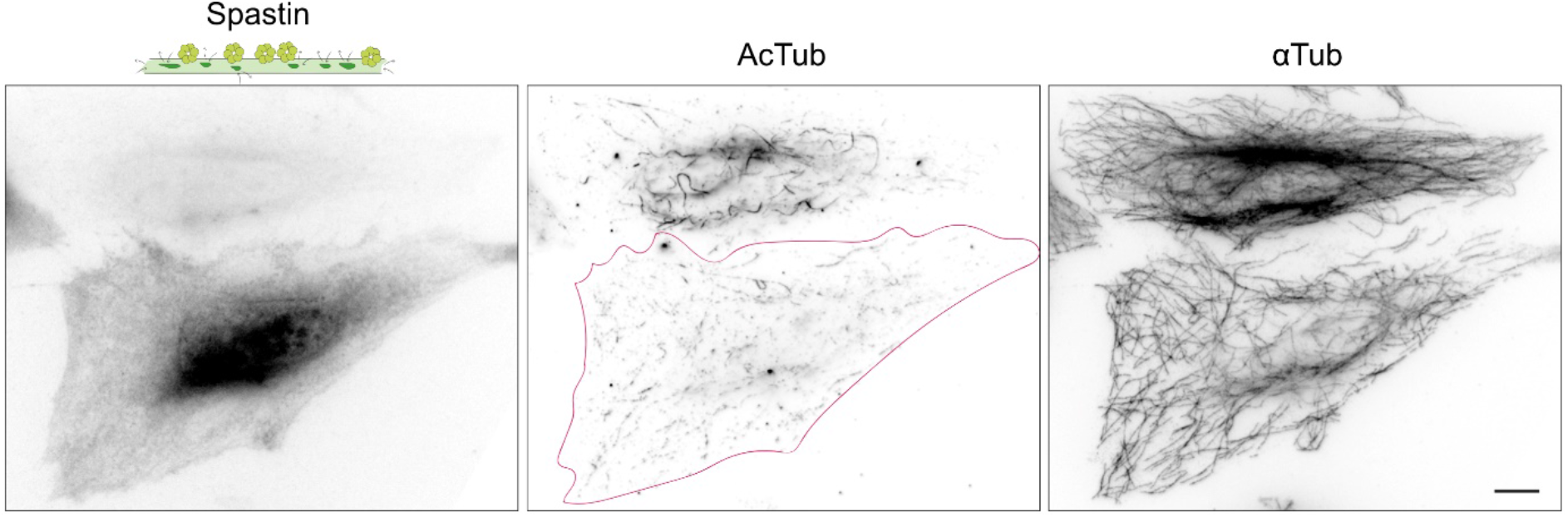
High levels of Spastin overexpression results in a fragmented microtubule network. Representative immunofluorescence of HeLa cells transfected with mCherry-Spastin. Cells were stained for AcTub and αTub. The magenta outline defines the edges of the cell overexpressing Spastin. Scale bar: 10 μm.

